# MuRF1 partners with TRIM72 to impair insulin signalling in skeletal muscle cells

**DOI:** 10.1101/2025.06.08.658491

**Authors:** Ibrahim Musa, Alex Peter Seabright, Yusuke Nishimura

**Affiliations:** School of Sport, Exercise and Rehabilitation Sciences, University of Birmingham, United Kingdom; Department of Human Physiology, Faculty of Basic Medical Sciences, College of Health Sciences, Prince Abubakar Audu University, Anyigba Kogi State, Nigeria; Research Institute for Sport & Exercise Sciences, Liverpool John Moores University, Liverpool, L3 3AF, United Kingdom

**Keywords:** protein-protein interaction, pulldown, E3 ligase, protein synthesis, glucose uptake

## Abstract

Muscle RING-finger protein 1 (MuRF1, gene name: *TRIM63*) is well known as a critical molecular regulator in skeletal muscle atrophy. Despite the identification of several substrates and interaction partners for MuRF1, the precise molecular mechanisms by which MuRF1 causes skeletal muscle atrophy remain unclear. To gain further insight into the underlying mechanism of skeletal muscle atrophy, we applied targeted biochemical approaches, and identified tripartite motif-containing protein 72 (TRIM72) as a novel MuRF1-interacting protein. Subsequent analysis using MuRF1 knockout and rescue experiments showed that TRIM72 protein abundance is dependent on the presence of MuRF1 protein. Furthermore, TRIM72 protein level was increased by dexamethasone treatment in C2C12 myotubes, alongside increased MuRF1 protein level. Dexamethasone decreases IRS1/Akt signalling, protein synthesis, and glucose uptake specifically in wild-type myotubes, but not in MuRF1 KO myotubes. Further analysis showed that overexpression of TRIM72 impairs IRS1/Akt signalling without the presence of MuRF1, indicating that MuRF1 induces a negative impact on insulin signalling through a plausible cooperation with TRIM72. Our findings provide novel non-degradative molecular roles of MuRF1 that link together skeletal muscle atrophy and impaired insulin responses.

**Highlights:**

- Identification of MuRF1 and TRIM72 interaction in skeletal muscle cells
- TRIM72 protein expression is dependent on the presence of MuRF1 protein
- Deletion of MuRF1 confers a protective effect against dexamethasone-induced impairment of IRS1/Akt signaling
- TRIM72 is sufficient to impair IRS1/Akt signaling

**Graphical Abstract:** 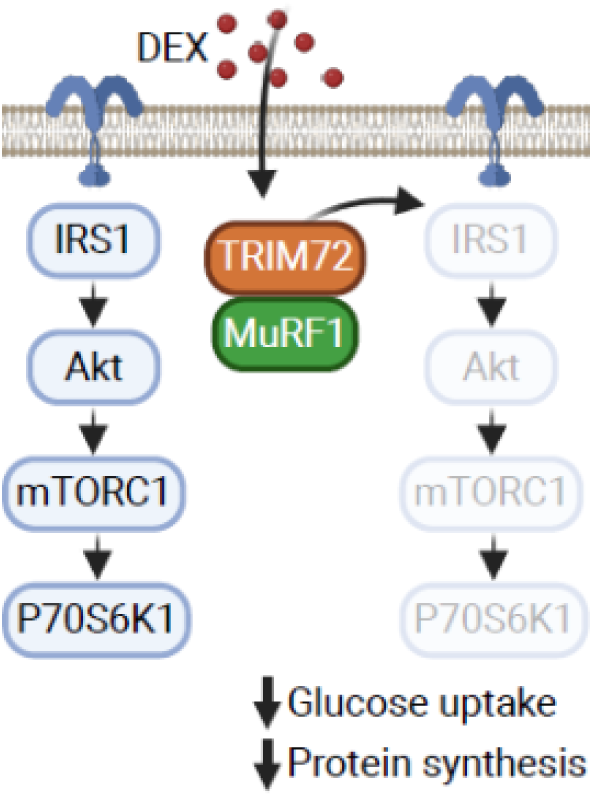

## Introduction

Muscle RING-finger protein 1 (MuRF1; gene name: *TRIM63*) is a muscle-specific E3 ligase highly critical in skeletal muscle atrophy [1, 2], but how MuRF1 regulates skeletal muscle mass at the molecular level remains poorly understood [3]. MuRF1 expression increases in several human and rodent models of denervation-, immobilization-, or unweighting-induced skeletal muscle atrophy [3–7]. Consistent with this, MuRF1 gene deletion can partially prevent denervation- and dexamethasone-induced skeletal muscle atrophy in mice [2, 3, 8]. Mechanistically, MuRF1 was reported to target myofibrillar proteins, such as myosin heavy chains (MHC), Myosin Binding Protein-C (MyBP-C), Myosin Light Chains 1 (MyLC1), and Myosin Light Chains 2 (MyLC2) for ubiquitylation and subsequent degradation via the ubiquitin proteasome system (UPS) to promote skeletal muscle atrophy [9, 10]. MuRF1 is widely accepted as a biomarker of muscle atrophy, but recent studies suggest that it may also regulate a range of other non-degradative cellular processes [8, 11, 12].

Impaired insulin signalling is known as a key driver of muscle protein degradation [13], primarily through reduced Akt activity, which leads to decreased FOXO phosphorylation, thereby increasing the expression of the E3 ubiquitin ligase MuRF1 [14, 15]. However, the molecular links between insulin signalling and MuRF1-dependent muscle atrophy remain unclear. A recent study led by Labeit *et al.,* [16] suggested that MuRF1 overexpression negatively regulate glucose metabolism in diabetic mice by impairing PI3K/Akt signalling. Their study showed that MuRF1 knockout (KO) mice exhibit an increased Akt phosphorylation at Ser473 in skeletal muscle [16]. Interestingly, treatment of MyoMed-205, a specific MuRF1 inhibitor stabilise the serum glucose in the diabetic mice [16]. Similarly, MuRF1 knockout mice revealed that MuRF1 negatively regulate insulin sensitivity [17]. These findings may point to a possible mechanistic links between MuRF1 and insulin signalling, raising the possibility that MuRF1 may play a role in insulin resistance and subsequent muscle wasting [18].

Besides MuRF1, several E3 ligases have been shown to regulate IRS1/ Akt signalling pathway. For example, Cbl-b (Casitas B-lineage lymphoma proto-oncogene-b), another RING-type E3 ligase catalyses the ubiquitylation of IRS1 protein and promotes its degradation in glucocorticoid induced atrophy [17]. Blocking the interaction between Cbl-b and the IRS1 protein using phosphopentapeptide DGpYMP, prevents glucocorticoid induced atrophy [19]. SCF-Fbxo40 is also another E3 ligase that ubiquitylates and degrades IRS1 upon IGF-1 stimulation [20]. In a previous study, SCF-Fbxo40 was found to be overexpressed in denervation-induced atrophy [21], whereas SCF-Fbxo40 knockdown in mice using siRNA (small-interfering RNA) results in thicker myotubes diameter [20]. TRIM72 (tripartite motif-containing protein 72), another muscle specific RING-type E3 ligase, ubiquitylates and promotes the degradation of IRS1 in cells [22, 23]. In contrast, TRIM72 knockdown in mice activates Akt and promotes myogenesis [23, 24]. While MuRF1, TRIM72, and Cbl-b E3 ligases have RING domains that facilitate ubiquitin transfer, MuRF1 and TRIM72 also have B-box and coiled-coil domains, that mediate protein-protein interactions [25]. Consistently, MuRF1’s B-box2 domain have been shown to interacts with muscle-type creatine kinase [12], and titin [26] in skeletal muscle. Surprisingly, a recent study demonstrated that MyoMed-205, a specific MuRF1 inhibitor, reduced both MuRF1 and TRIM72 levels in obese ZSF1 rats [27], indicating a possible functional interaction between these two E3 ligases.

The number of substrates that MuRF1 targets for ubiquitylation is currently unclear, making it difficult to understand how MuRF1 overexpression causes muscle atrophy. A recent study led by Baehr *et al.,* [2], suggests that MuRF1 could induce muscle atrophy via previously unexplored non-degradative mechanisms. In their study, MuRF1 overexpression in mice was sufficient to cause atrophy, but the majority of its ubiquitylated target proteins do not undergo degradation [2]. Further analysis by Baehr *et al.,* [2] showed that MuRF2, MuRF3, and TRIM25 E3 protein abundances were positively correlated with MuRF1 overexpression [2], demonstrating that MuRF1 may coordinate with other E3 ligases to regulate skeletal muscle mass. This highlights the need to explore MuRF1’s non-canonical roles and its interactors.

This study examined the molecular function of MuRF1 and its interplay with TRIM72 in skeletal muscle myotubes. We identified TRIM72 as a novel interacting partner of MuRF1 and validated their association in living cells. Skeletal muscle myotubes lacking MuRF1 showed a significant reduction in TRIM72 protein, while MuRF1 rescue experiment restored TRIM72 protein abundance. To examine the functional significance of this interaction, we compared wild-type and MuRF1 knockout myotubes under dexamethasone (Dex)-induced atrophy conditions. Dexamethasone treatment upregulated both MuRF1 and TRIM72, concomitant with impaired IRS1/Akt signaling, reduced protein synthesis, and diminished glucose uptake, effects that were absent in MuRF1-deficient cells. These findings establish TRIM72 as a critical effector of MuRF1’s role in muscle atrophy and insulin resistance, providing new mechanistic insights into the interplay between proteolytic and metabolic pathways in skeletal muscle.

## Results

### Identification of TRIM72 as a novel MuRF1-interacting protein

The convergence of TRIM72 and MuRF1, on IRS1 degradation and the finding that MuRF1 inhibition decreases TRIM72 prompted us to firstly investigate the interactions between these two proteins. To investigate this, we first generated L6 skeletal muscle cell lines that stably express GFP-MuRF1 or GFP alone, as a control. The generated cells were differentiated into myotubes. Protein lysates were prepared and subjected to GFP pulldown, followed by immunoblotting for TRIM72. Our result showed that TRIM72 co-immunoprecipitated with MuRF1, confirming a direct interaction between these two proteins in cell (Fig 1A, upper panel).

**Figure 1.**
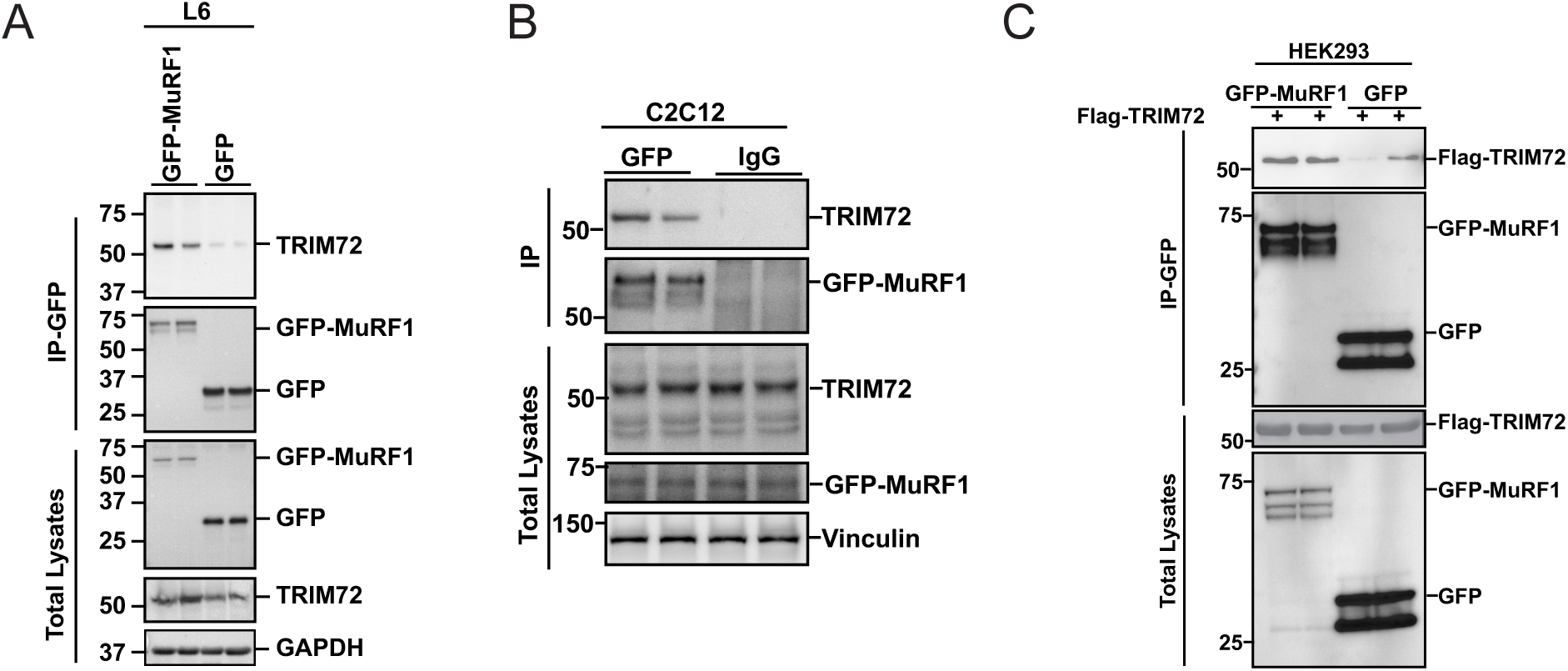
MuRF1 interacts with TRIM72 in skeletal muscle cells. (A) Verification of MuRF1 interaction with endogenous TRIM72 in L6 myotubes that stably expressed GFP-MuRF1. Total lysates (1 mg) from L6 myotubes that stably expressed GFP-MuRF1 or GFP-empty were subjected to GFP pulldown using GFP-Trap agarose beads. The pulldown products were immunoblotted with the indicated antibodies. To confirm protein expression and loading control, total lysates (lower panel) were immunoblotted with the indicated antibodies. Immunoblots were representative of three independent experiments. (B) Verification of endogenous MuRF1 interaction with endogenous TRIM72 in GFP-MuRF1 knock-in C2C12 myotubes. Total lysates (1 mg) from GFP-MuRF1 knock-in C2C12 myotubes were subjected to GFP pulldown using GFP-Trap agarose beads or beads bound with IgG (as a control). The pulldown products and immunoprecipitates were immunoblotted with the indicated antibodies. To confirm protein expression and loading control, total lysates (lower panel) were immunoblotted with the indicated antibodies. Immunoblots were representative of three independent experiments. (C) Verification of MuRF1 and TRIM72 interaction in HEK293 cells. HEK293 cells were transiently co-transfected with FLAG-TRIM72 and either with GFP-MuRF1 or GFP-empty (as a control) for 48 h. Total lysates (0.5 mg) was subject to pulldown using GFP-Trap agarose beads. The pulldown products were immunoblotted with the indicated antibodies. To confirm exogenous protein expression, total lysates (lower panel) were immunoblotted with the indicated antibodies. Immunoblots were representative of one experiment.

Due to the lack of antibodies that are appropriate for MuRF1 immunoprecipitation, MuRF1 N-terminal GFP knock-in (GFP-MuRF1) C2C12 muscle cells were generated using CRISPR/Cas9 technology to test the interaction between MuRF1 and TRIM72 under endogenous condition and in a different muscle cell line. Again, GFP pulldown showed that TRIM72 co-immunoprecipitated with GFP-MuRF1, but not with IgG (Fig 1B). These results demonstrated that MuRF1 and TRIM72 physically interact also in C2C12 skeletal muscle cells. To further confirm the interaction between MuRF1 and TRIM72 in another cell type, we transiently co-transfected Flag-TRIM72 either with GFP-MuRF1 or GFP-empty in HEK293 cells. The GFP pulldown products were immunoblotted with the Flag-tagged HRP conjugated antibody for TRIM72. The results showed that TRIM72 was pulled down together with GFP-MuRF1, and was dramatically reduced in GFP alone (Fig 1C, upper panel), confirming again the interaction between MuRF1 and TRIM72 in different cells. These results suggest that MuRF1 may stabilize or promote TRIM72 expression explaining why MuRF1 inhibition reduces TRIM72 protein levels in a previous study [27].

### TRIM72 protein expression is dependent on the presence of MuRF1 protein in C2C12

We next examined whether TRIM72 constitutes a functional interaction partner or represents a substrate of MuRF1 protein. We used CRISPR/Cas9 technology to generate a MuRF1 knockout (KO) in C2C12 skeletal muscle cell line. Myoblasts of wild-type (WT) or two different clones of MuRF1 KO C2C12 were differentiated into myotubes. As expected, MuRF1 protein content was not detected in MuRF1 KO cells, indicating the successful knock out of MuRF1 (Fig 2A). Notably, TRIM72 protein levels were significantly reduced in MuRF1 KO myotubes, compared to WT (Fig 2A and 2B). We next examined if a transient reintroduction of MuRF1 could restore TRIM72 expression. TRIM72 protein expression was partially restored when Flag-MuRF1 was transiently overexpressed in the MuRF1 KO cells, (Fig 2C and 2D). It is known from previous studies that these two proteins increase during the differentiation [24, 28]. Thus, the restoration of TRIM72 expression when MuRF1 was added in MuRF1 KO myotubes led us to compare their expression during the differentiation time course (Fig 2E and 2F). The data showed that TRIM72 protein expression is associated with MuRF1 expression along the differentiation time course, but MuRF1 protein expression precedes TRIM72 protein expression (Fig 2E and 2F). Taken together, our results suggest that the expression of TRIM72 protein is MuRF1 expression dependent in skeletal muscle cells. Previous studies demonstrated that TRIM72 is a known negative regulator of IRS1/Akt signalling [22, 29]. We therefore further explored the roles of MuRF1 and TRIM72 in IRS1/Akt signalling.

**Figure 2.**
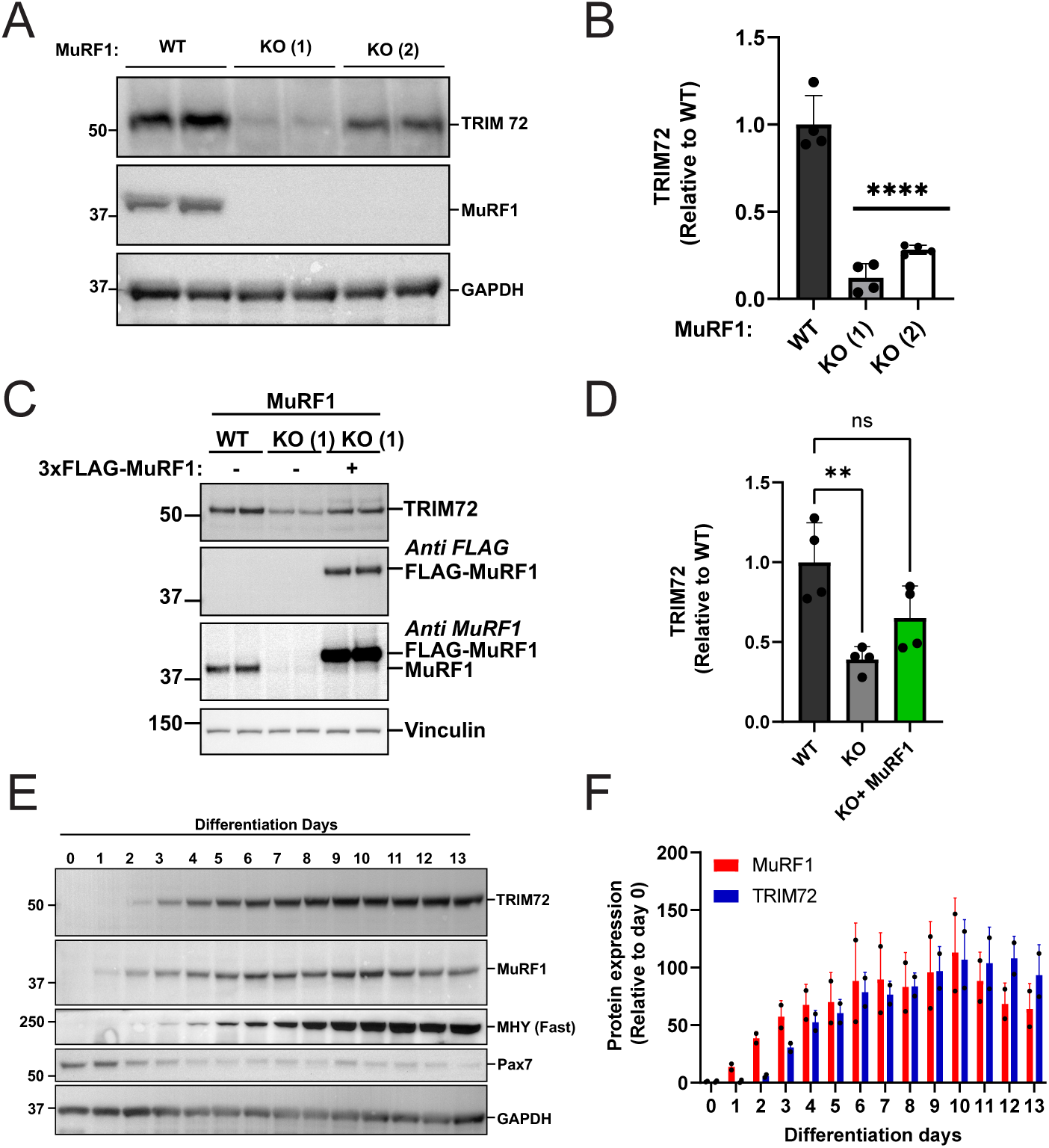
TRIM72 expression is associated with MuRF1 expression in C2C12 myotubes. (A) TRIM72 protein levels are reduced in MuRF1 knockout C2C12 myotubes. Total lysates from WT (wild type) or two different MuRF1 knockout (KO 1 & 2) C2C12 myotubes were analysed by immunoblotting and probed with the indicated antibodies. Immunoblots were representative of four independent experiments (B) Quantification of TRIM72 protein expression in WT and MuRF1 KO C2C12 myotubes. Quantitative data from (A) were normalized to WT and subjected to one-way analysis of variance (ANOVA) before Dunnett’s multiple comparisons post-hoc test analysis. Error bars indicate the mean ± standard deviation (n = 4). **** p < 0.0001 compared to WT. (C) Transient overexpression of MuRF1 restored TRIM72 protein expression in MuRF1 KO myotubes. Total lysates of C2C12 myotubes from WT or MuRF1 KO with and without transient transfection of 3xFLAG-MuRF1 for 48 h were analysed by immunoblotting and probed with the indicated antibodies. Immunoblots were representative of four independent experiments. (D) Quantification of TRIM72 protein expression in WT and MuRF1 KO with and without transient transfection of 3xFLAG-MuRF1. Quantitative data from (C) were normalized to WT and subjected to one-way analysis of variance (ANOVA) before Dunnett’s multiple comparisons post-hoc test analysis. Error bars indicate the mean ± standard deviation (n = 4). **** p < 0.0001 compared to WT. (E) Representative blots of two independent experiments showing MuRF1 and TRIM72 protein expression during differentiation time course for up to 13 days in C2C12 cells. Total lysates from C2C12 cells at the indicated days of differentiation were analysed by immunoblotting and probed with the indicated antibodies. (F) Bar chart analysis of MuRF1 and TRIM72 protein expressions during differentiation time course in C2C12 cells. Quantitative data from (E) were normalised to day 0 and presented as bar chart. n = 2 in each time point.

### Dexamethasone increases TRIM72 and MuRF1 protein expressions while decreasing IRS1/Akt signalling in C2C12 myotubes

MuRF1 protein content is known to be increased in dexamethasone (Dex) treated C2C12 myotubes [30]. We next examined whether TRIM72 protein is similarly upregulated in dexamethasone treated C2C12 myotubes. As expected, Dex treatment increased MuRF1 and also TRIM72 protein expression (Fig 3B & 3C). Pearson’s correlation coefficient analysis from the Dex dose-response treatment data (Fig 3D) shows a positively correlation between TRIM72 and MuRF1 protein expressions (r = 0.698, *P* < 0.0001). Previous study showed that overexpression of TRIM72 reduces IRS1 protein levels in C2C12 myotubes [22]. After showing that MuRF1 and TRIM72 proteins are both upregulated after Dex treatment, we wanted to know whether IRS1 protein level is also reduced by Dex treatment. We observed that IRS1 protein level (Fig 3E), Akt phosphorylation at Ser 473 (Fig 3F) and Thr 308 (Fig 3G) were reduced by 1 μM of Dex treatment where TRIM72 and MuRF1 protein content increased. Additionally, the reduced phosphorylation status in the downstream of mTOR, such as p70 S6K1 phosphorylation at Thr 389, further supported the decreased Akt activity (Fig 3H). Taken together, these results show that MuRF1 and TRIM72 proteins are both upregulated by Dex treatment with a concomitant reduction of the IRS1/Akt signalling. Based on the effectiveness of the Dex concentrations (Figures 3A & 3C), 1 μM Dex was chosen for the subsequent experiments.

**Figure 3.**
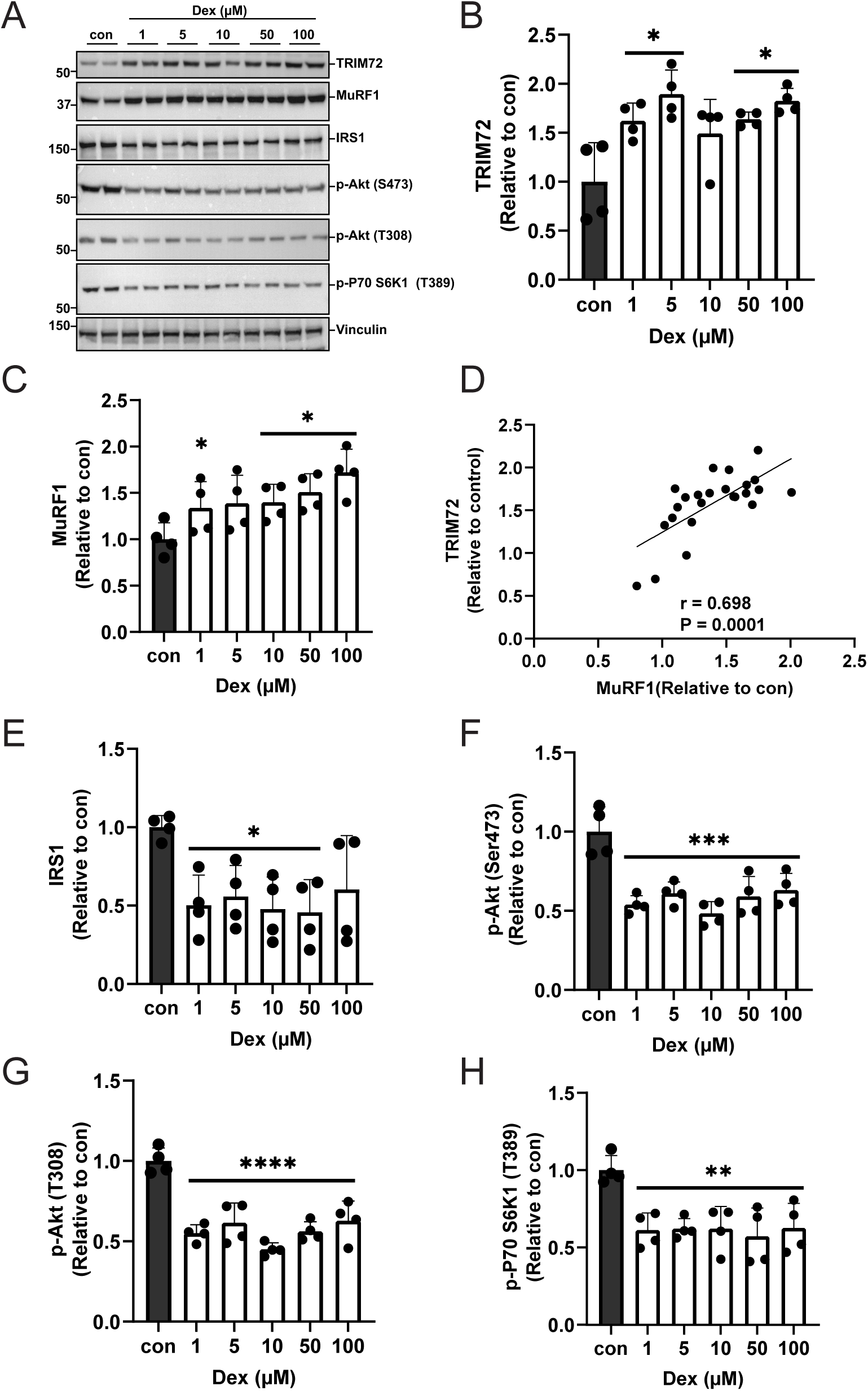
Dexamethasone increases TRIM72 and MuRF1 protein expression while decreasing proximal insulin signalling in C2C12 myotubes. Total lysates from C2C12 myotubes treated with 0.1% ethanol as controls (con) or with indicated concentrations of dexamethasone (Dex) for 24 h were analysed by immunoblotting and probed with the indicated antibodies. (A) Representative blots showing the effect of dexamethasone on TRIM72, MuRF1 proteins, and proximal insulin signalling in C2C12 myotubes. Immunoblots were representative of four independent experiments. (B, C, E-H) Quantitative data of TRIM72, MuRF1 and IRS1 protein expressions and proximal insulin signalling, including Akt and p70 S6K1 phosphorylation status. Quantitative data were normalised to control (con) and subjected to one-way analysis of variance (ANOVA) before Dunnett’s multiple comparisons post-hoc test analysis. Error bars indicate the mean ± standard deviation (*n* = 4 independent experiments). Statistical analysis was considered significant at * *p* < 0.05, ** *p* < 0.01, *** *p* < 0.001, **** *p* < 0.0001, compared with control. (D) TRIM72 expression is associated with MuRF1 protein expression in C2C12 myotubes treated with dexamethasone. Quantitative data of TRIM72 and MuRF1 from *n* = 4 independent experiments were subjected to Pearson’s correlation coefficient analysis to examine their protein expression correlation.

### Dexamethasone-induced downregulation of proximal insulin signalling, insulin-stimulated protein synthesis and glucose uptake are prevented in MuRF1 KO C2C12 myotubes

MuRF1 knockout cells were used to further confirm whether the TRIM72 mediated downregulation of IRS1/Akt signalling is MuRF1 dependent. The results showed that Dex treatment reduced IRS1 protein levels and Akt phosphorylation at serine 473 in WT, but not in MuRF1 KO myotubes (Fig 4A). These data confirm that MuRF1 expression is required for the TRIM72-mediated downregulation of IRS1/Akt signalling in Dex treated C2C12 myotubes. To further confirm that MuRF1 is required for Dex-mediated global insulin responsiveness, we measured protein synthesis and glucose uptake. Dex treatment significantly reduced protein synthesis in WT, but this reduction is absent in MuRF1 KO myotubes (Fig 4F and 4G). Similar results were found in insulin-stimulated 2-Deoxyglucose (2DG) uptake, where Dex treatment reduced glucose uptake in WT, but not in myotubes of MuRF1 KO (Fig 4H). These results suggest that MuRF1 is required for TRIM72-mediated reduction in IRS1/Akt signalling, which in turn, alters insulin-mediated metabolism, including protein synthesis and glucose uptake.

**Figure 4.**
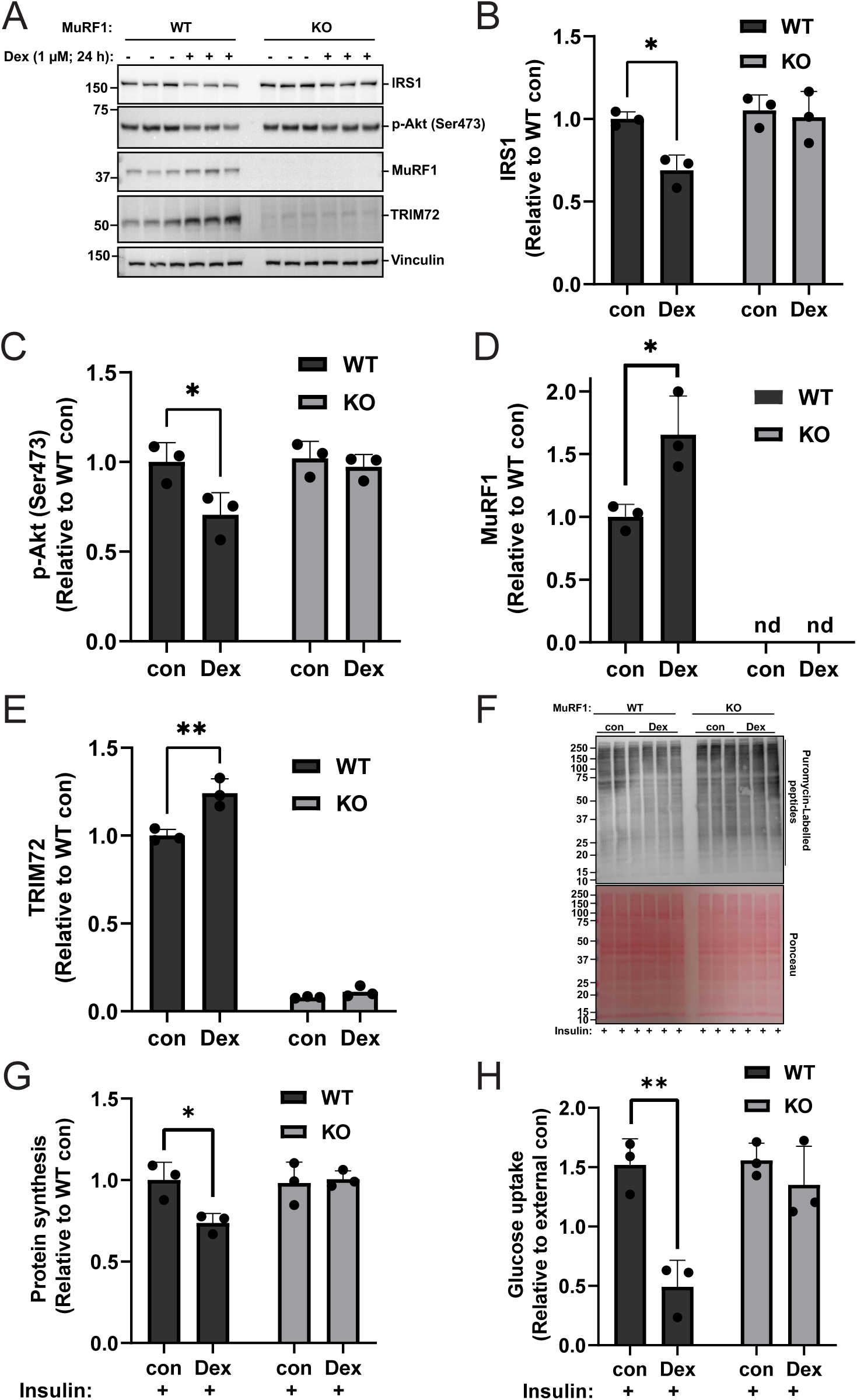
MuRF1 KO prevents dexamethasone-induced downregulation of proximal insulin signalling, insulin-stimulated protein synthesis and glucose uptake in C2C12. (A) Immunoblots from three independent experiments showing IRS1, p-Akt (Ser473), MuRF1 and TRIM72 in WT and MuRF1 KO C2C12 myotubes treated with and without dexamethasone. Total lysates from WT or MuRF1 KO C2C12 myotubes treated with and without 1 µM dexamethasone (Dex) for 24 hours were analysed by immunoblotting before probing with the appropriate antibodies. (B-E) Figures showing quantitative data of IRS1 protein expression (B), Akt Ser473 phosphorylation (C), MuRF1 (D) and TRIM72 (E) protein expressions. Quantitative data from (A) were normalised to wild-type control (WT con) and subjected to a two-way ANOVA before Tukey’s multiple comparisons post-hoc test analysis. Error bars indicate the mean ± standard deviation (*n* = 3). \**p* < 0.05, \*\**p* < 0.01, compared to WT con. nd: non-detectable. (F) Immunoblot showing insulin-stimulated protein synthesis and loading control (ponceau) in WT or MuRF1 KO C2C12 myotubes treated with and without dexamethasone. WT or MuRF1 KO C2C12 myotubes were pre-treated with 1 µM puromycin and either 0.1% ethanol (control), or 1 µM dexamethasone for 24 h before stimulating by 100 nM insulin for 30 minutes prior to cell lysis in a serum free media. The protein synthesis (incorporated puromycin-labelled peptides) was determined by immunoblotting with anti-puromycin antibody. (G) Quantification of insulin-stimulated protein synthesis in WT or MuRF1 KO myotubes. Quantitative data from (F) were normalised to wild-type control (WT con) and subjected to a two-way ANOVA before Tukey’s multiple comparisons post-hoc test analysis. Error bars indicate the mean ± standard deviation (*n* = 3 independent experiments). \**p* < 0.05, compared to WT con. (H) Insulin-stimulated 2-Deoxyglucose (2DG) uptake in WT and MuRF1 KO treated with or without dexamethasone. C2C12 WT and MuRF1 KO myotubes pre-treated with or without 1 µM dexamethasone for 24 hours were serum starved overnight, stimulated by 1 µM insulin for 1 h before measuring 2DG uptake for 30 min. Results were presented as glucose uptake (2DG fold-changes) relative to external controls (without insulin stimulation). Data were analysed by a two-way ANOVA before Tukey’s multiple comparisons post-hoc test analysis. Error bars indicate the mean ± standard deviation (*n* = 3 independent experiments). \*\**p* < 0.01 compared with insulin-stimulated control.

### Restoration of TRIM72 in C2C12 MuRF1 KO myotubes decreases proximal insulin signalling

To verify if TRIM72 is an essential intermediary between MuRF1 and Dex-induced reduction in IRS1/Akt signalling, we conducted a rescue experiment by transiently transfected Flag-TRIM72 into MuRF1 KO myoblasts. As expected, overexpression of TRIM72 in MuRF1 KO significantly reduced IRS1 protein (Fig 5A and 5B) and phosphorylation of its downstream target, such as Akt at Ser 473 and p70 S6K1 at Thr 389 (Fig 5). Overall, our data demonstrated that restoration of TRIM72 in MuRF1 KO is sufficient to reduce IRS1 protein and its downstream signalling targets by Dex treatment. This observation indicates that TRIM72 is sufficient to impair IRS1/Akt signalling, but TRIM72 protein is controlled by a MuRF1-dependent manner, highlighting both MuRF1 and TRIM72 as key regulators for IRS1/Akt signalling in skeletal muscle cells.

**Figure 5.**
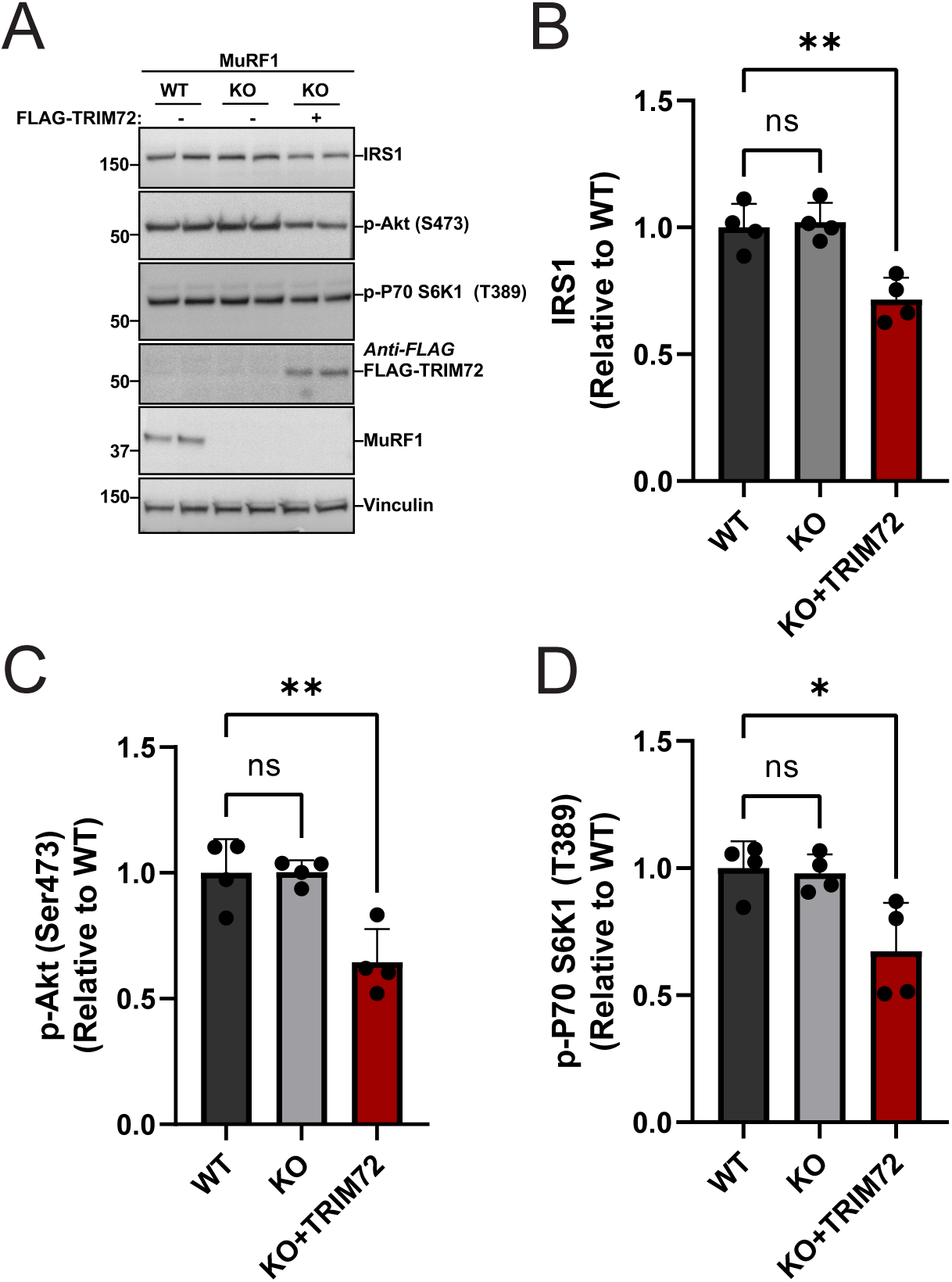
Restoration of TRIM72 in C2C12 MuRF1 KO myotubes decreases proximal insulin signalling. (A) Immunoblots from four independent experiments showing IRS1, p-Akt (Ser473), p70 S6K1 (T389), TRIM72 and MuRF1 in WT and MuRF1 KO C2C12 myotubes. Total lysates of C2C12 myotubes from WT or MuRF1 KO with and without transient transfection of FLAG-TRIM72 for 48 hours were analysed by immunoblotting and probed with the appropriate antibodies. (B-D) Figures showing quantitative data of IRS1 protein expression (B), Akt Ser (473) phosphorylation (C) and p70 S6K1 (T389) phosphorylation (D). Quantitative data from (A) were normalised to wild-type (WT) and subjected to a one-way ANOVA before Dunnett’s multiple comparisons post-hoc test analysis. Error bars indicating: mean ± standard deviation (*n* = 4 independent experiments). \**p* < 0.05, \*\**p* < 0.01, compared to Wild Type. ns: non-significant.

## Discussion

In this present study, we discovered TRIM72 as a novel MuRF1 effector protein in skeletal muscle cells. Our findings demonstrated that MuRF1 stabilises TRIM72, enabling TRIM72 to downregulate IRS1/Akt signalling and impair insulin-stimulated protein synthesis and glucose uptake under dexamethasone treatment in skeletal muscle cells. MuRF1 KO myotubes further confirm the requirement of MuRF1 for the detrimental effects of TRIM72 under dexamethasone treatment though TRIM72 itself is sufficient to impair IRS1/Akt signalling. These results suggest that MuRF1 may regulate muscle mass by acting as a negative regulator of anabolic pathways. Thus, this study provides new evidence that MuRF1 can influence skeletal muscle homeostasis through non-degradative pathway, highlighting the need for additional exploration of MuRF1’s molecular roles.

It is interesting to note that TRIM72 protein abundance was reduced in MuRF1 knockout myotubes (KO) but was restored when MuRF1 was added (Fig 2). This suggests that TRIM72 is not a MuRF1 substrate but rather a partner whose protein expression is regulated by MuRF1. Consistently, differentiating C2C12 cells start to express MuRF1 proteins before TRIM72 proteins over a 13-day period of differentiation (Fig 2), supporting our notion that MuRF1 regulates the levels and stability of TRIM72 protein in skeletal muscle. In agreement with this, TRIM72 was downregulated in *tibialis anterior* (TA) muscle after MuRF1 inhibitor (MyoMed 205) was administered to obese ZSF1 rats in a previous study [27]. This finding supports our hypothesis that MuRF1 interacts with and modulates TRIM72.

Here we showed that MuRF1 and TRIM72 proteins are both upregulated after Dex treatment. In a common model of skeletal muscle atrophy, dexamethasone (Dex) treatment is well known to cause atrophy in part through increasing MuRF1 levels [8, 9, 17, 30, 31]. For the first time, we showed Dex treatment upregulates TRIM72 protein abundance (Fig 3B). The upregulation of TRIM72 protein in Dex treated C2C12 myotubes, corresponded with a dramatic decreased in IRS1 protein levels, corroborating previous findings that TRIM72 facilitates IRS1 degradation [22]. Several studies have reported that TRIM72 ubiquitylates and degrades IRS1 [22–24] via the 26S proteasome, leading to impaired IRS1 signalling [Fig. 3]. Likewise, the overexpression of MuRF1 was demonstrated to reduce IRS1 levels [17], underscoring the shared function of these E3 ligases in inhibiting insulin signalling to promote muscle atrophy. This is consistent with a previous study that IRS1 deficiency promote muscle atrophy both *in vivo* and *in vitro* [32, 33]. Thus, our data provide a potential mechanistic link between muscle wasting and insulin resistance in skeletal muscle.

IRS1/Akt/mTORC1 pathway is known to promote protein synthesis [34–37]. MuRF1 has been shown to decrease protein synthesis under conditions of nutrient deprivation model of atrophy [12]. MuRF1 overexpression reduces IRS1 protein levels [17], which may explain the impairment of protein synthesis via IRS1/Akt signalling. However, the underlying molecular mechanisms that control this process was yet to be uncovered. Our findings show that MuRF1 and its interaction partner, TRIM72, negatively affect insulin-induced protein synthesis after dexamethasone treatment (Fig 4F). In skeletal muscle, TRIM72 functions as an autonomous ubiquitin ligase to degrade the IRS1 protein as a negative regulator of protein synthesis [22–24]. The increase in the insulin-induced protein synthesis observed in MuRF1 KO C2C12 myotubes (Fig 4F), could be attributed to the restored insulin signalling activity caused by the marked loss of TRIM72 in the MuRF1 KO C2C12 myotubes. This finding is consistent with a previous study where MuRF1 KO was demonstrated to be resistant to reductions in protein synthesis induced by dexamethasone [8].

Glucose metabolism is essential for maintaining organ function in humans [38]. Dex treatment significantly reduced insulin-induced glucose uptake in WT but not in MuRF1 KO myotubes when compared with controls (Fig. 4H). Recently, Labeit *et al.,* [16] examined the effect of MuRF1 on glucose metabolism in diabetic mice. They found diabetic mouse exhibit an impaired PI3K/Akt signalling, but MuRF1 KO mice exhibit an increased Akt phosphorylation at Ser473 in skeletal muscle [16]. Consistently, MuRF1 inhibitor (MyoMed-205) was shown to stabilise serum glucose concentration in diabetic mice [16]. In line with our findings, previous studies [22, 39, 40] also reported that TRIM72 is sufficient to promote diabetes, via downregulation of IRS1. However, a contrasting role of TRIM72 in the regulation of diabetes has also been reported [41, 42]. These studies showed that neither the injection of recombinant human TRIM72 (rhTRIM72) nor the adenoviral gene transfer of TRIM72 was able to cause changes in blood glucose in either diabetic db/db or nondiabetic mice [41, 42]. Interestingly, the disparity in these studies revealed that the negative effects of rhTRIM72 on metabolism are only seen in cases of advanced diabetes [40], emphasizing the need to examine E3 ligases in context-specific disease models.

## Conclusion

Our study identifies the MuRF1-TRIM72 axis as a crucial regulatory mechanism in dexamethasone-induced metabolic dysfunction. Our findings suggested that insulin-stimulated protein synthesis and glucose uptake in skeletal muscle are impaired by a MuRF1-dependent regulation of TRIM72 protein levels. This finding reveals a previously unknown function of MuRF1 in non-proteolytic metabolic regulation and offers a new perspective on the molecular mechanism underlying insulin resistance and muscle atrophy. We propose a mechanistic model wherein the upregulation of MuRF1 raises and stabilises the levels of TRIM72 protein, which in turn impairs the anabolic processes triggered by insulin.

## Materials and methods

### Antibodies and Reagents

The following antibodies were applied for western blot analysis: Anti-MuRF1 (Santa Cruz SC-398608; 1:1000), Anti-TRIM72 (Antibodies.com A84884; 1:8000), Anti-phospho Akt (Ser473) (Cell Signalling Technology 4060; 1:1000), Anti-phospho-Akt (T308) (Cell Signalling Technology 2965; 1:1000), Anti-phospho-P70 S6K1 (T389) (Cell Signalling Technology 9234; 1:1000), Anti-GAPDH (Cell Signalling Technology 5174; 1:1000), Anti IRS1 (Cell Signalling Technology 3407; 1:1000), Anti-GFP (Chromotek 3H9-100; 1:2000), Anti-FLAG M2 (Sigma-Aldrich F1804; 1:1000), Anti-Vinculin (Abcam ab73412; 1:1000), and Anti-puromycin (Sigma-Aldrich P8833; 1:1000). Anti-Myosin (fast) (Sigma M4276; 1:1000), Anti-PAX7 antibody (ThermoFisher; PA1-117, 1:1000), and Anti-Myosin (Slow) (Sigma M8421; 1:1000). The following secondary HRP-linked antibodies were applied: Anti-goat (Cell Signalling Technology 7077; 1:10 000), Anti-mouse (Cell Signalling Technology 7076; 1:10 000), Anti-rabbit (Cell Signalling Technology 7074; 1:10 000), and Anti-rat (Cell Signalling Technology 7077; 1:5000) Antibodies. The reagents for cell culture that were used includes: High glucose GlutaMAX Dulbecco’s Modified Eagle Medium with 1 mM of sodium pyruvate (Thermo Fisher Scientific, Loughborough, UK, 31966021); Cytiva Hyclone Foetal bovine serum (Fisher Scientific, South America, 10309133); Penicillin-Streptomycin (10 000 Units/mL-ug/mL); Horse serum (Sigma-Aldrich, Cambridgeshire, UK, H1270); DPBS (Sigma, 14190); polybrene infection reagent (Merck life scientific UK, TR-1003); and 0.05% Phenol red Trypsin-EDTA (Fisher Scientific, 25300062); Polyethylenimine (PEI) solution (Sigma, 408727); GAG/POL and VSV-G plasmids purchased from Clonetech (Saint-Germain-en-Laye, France). Chemicals/compounds such as Dexamethasone (Sigma-Aldrich D4902) and Insulin solution human (Sigma-Aldrich, Poole, UK; I9278) were used.

### Cell lines and Culture

L6 rat and C2C12 mouse skeletal muscle myoblasts were purchased from the American Type Culture Collection (ATCC, Manassas, VA, USA). GFP-MuRF1 knocked-in (KI) and MuRF1 knockout (KO) C2C12 cell lines were developed using CRISPR/Cas9 system as was described previously [43].

Firstly, the sense and antisense sgRNA constructs for A (tCTGATTCCTGATGGAAACGCTA, GCTGATCTGCCCCATCTGCCTtG); sense and antisense sgRNA constructs for B (tCTGGAGAAGCAGCTGATCTGCC, tGAGATGTTTACCAAGCCTGTcG), which target the N-terminal GFP knock-in to the MuRF1 locus, and the Nter GFP donor (pMK-RQ vector, DU60520) were generated via the CRISPR vector designing tool (http://tools.genome-engineering.org). The generated oligonucleotides identified were annealed to their respective complements with the cloning tag ‘a,’ ‘g’ as was shown in the following: Nter KI as A (TAGCGTTTCCATCAGGAATCAGa, CaAGGCAGATGGGGCAGATCAGC) and Nter KI as B (GGCAGATCAGCTGCTTCTCCAAGa, CgACAGGCTTGGTAAACATCTCa) to generate dsDNA inserts with compatible over-hangs to BbsI-restriction site of pBabeD-puro and pX335-Cas9-D10A vectors. The antisense sgRNA was cloned onto pX335-spCas9-D10A (Addgene, 42335) and the sense sgRNA cloned onto the pBabeD-puro (puromycin selectable plasmid P U6). C2C12 myoblasts cells of 60-70% confluency was co-transfected with 1 µg CRISPR plasmids and 3 µg of the fluorescent GFP tag donor plasmid using polyethylenimine (PEI) transfection reagent. After 48 h co-transfection, the transfected C2C12 mouse skeletal muscle cell was selected using 2 μg/mL puromycin. The selected pool of clones was subsequently single-cell sorted and collected into 96 well plates using the ARIA fusion sorter (BD Biosciences, Berkshire, UK). This was carried out by a specialist at the University of Birmingham’s flow cytometry center (Institute of Biomedical Research). Positive clones for GFP-MuRF1 KI and MuRF1 KO C2C12 were validated by immunoblotting.

L6 stably expressing GFP-MuRF1 and GFP empty were generated using a retrovirus encoding human MuRF1 protein fused with a GFP tag at the N-terminus or a retrovirus encoding a GFP empty respectively. Briefly, cDNA plasmids for a human MuRF1 protein fused with a GFP tag at the N-terminus or a GFP empty were cloned into a pBABED.puro vectors respectively. The construct was co-transfected into HEK293 FT cells with GAG/POL and VSV-G expression plasmids (Clonetech, Saint-Germain-en-Laye, France) for retrovirus production using polyethylenimine (PEI) transfection reagent in accordance with manufacturer’s instructions. Virus was harvested 48 hours after transfection and applied to L6 skeletal muscle myoblasts in the presence of 10 μg/mL polybrene (Sigma-Aldrich, Cambridgeshire, UK). After 48 h infection, the infected L6 rat skeletal muscle cell was selected using 2 μg/mL puromycin. Positive GFP-MuRF1 and GFP were cultured for further expansion and cryopreserved after being validated by western blotting. All myoblasts cells were cultured and grown in high-glucose Dulbecco’s Modified Eagle Medium (DMEM) supplemented with 10% (v/v) foetal bovine serum and 1% (v/v) Penicillin-Streptomycin (10 000 units/mL-μg/mL). At about 90% confluency, skeletal muscle myoblasts were differentiated into myotubes for at least five days using high-glucose DMEM supplemented with 2% (v/v) horse serum (HS) and 1% (v/v) penicillin-streptomycin (10 000 units/mL-μg/mL). To maintain cells or myotubes viability, the medium was changed every two days until cells were fully differentiated. Description of drug treatment on myotubes were discussed in the respectively figure legends.

### Transient transfection of mammalian cell lines

Transient transfections were performed using Polyethylenimine (PEI) transfection reagent. Specifically, maintained culture cells or myotubes were transfected with the required plasmid DNA in a 1:2.5 ratio (µl/µg) of PEI to the plasmid. Briefly, PEI and plasmid DNA were firstly mixed individually, in a separate eppendorf tubes containing 500 µl DMEM devoid of serum for 5 min at room temperature. After 5 min incubation, the PEI/DNA mixture was combined into one and was further incubated for 20 min before added dropwise onto the cells or myotubes. After overnight incubation, fresh media were replaced to wash out PEI remnant after transfection. At 48 h post transfection, cells were lysed and harvested for analysis.

### Cell lysis

At the end of the experiments, the prewashed cells or myotubes with ice-cold DPBS were lysed and collected in cold sucrose lysis buffer containing 250 mM of sucrose, 10 mM of sodium β-glycerolphosphate, 50 mM of Tris-base (pH 7.5), 5 mM of sodium pyrophosphate, 50 mM of sodium fluoride, 1 mM of EDTA, 1 mM of benzamidine, 1 mM of EGTA, 1 mM of sodium orthovanodate, 1% of Triton X-100, 1 x complete Mini EDTA-free protease inhibitor cocktail, and 100 mM of 2-chloroacetamide. Lysates were pelleted for 15 minutes at 4°C at 13,000 rpm, and the supernatant was collected and stored at −80°C for analysis. Bradford protein quantification assay was performed on the protein extracts with BSA as protein standards.

### Sample preparation

Protein lysates were prepared using 1x LDS sample buffer (NuPAGE, Invitrogen, NP0008). Samples were left overnight in 1.5% of 2-mercaptoethanol to denature at room temperature before analysis.

### GFP pull down

20 μl of GFP-trap agarose or IgG bead slurry pre-washed with an ice-cold DPBS was then washed twice with ice-cold sucrose lysis buffer before incubated with 2 mg lysates overnight on a rotating wheel at 4 °C. Beads were then pelleted by centrifugation at 3500 rpm for 1 min at 4°C. After pelleted (washing) three times with sucrose lysis buffer, the co-immunoprecipitated proteins were eluted with 2x NuPAGE LDS sample buffer. Samples were left overnight in 1.5% of 2-mercaptoethanol to denature at room temperature before analysis.

### Western blotting

Aliquots from the prepared samples were loaded and run on 10% Bis/Tris gels (ThermoFisher Scientific, Leicestershire, UK) in 1x MOPS buffer (ThermoFisher Scientific, Leicestershire, UK) for approximately 75 minutes at 140 V. Proteins from the gel were transferred to 0.2 μM PVDF (polyvinylidene fluoride) membranes (Millipore, Hertfordshire, UK) in 1x transfer buffer containing; 20 mM Tris base, 150 mM glycine, and 20% methanol for 1 h at 100 V and 4°C condition. The membranes were blocked for 1 hour in 5% (w/v) dried skimmed milk diluted with TBS-T (Tris-buffered saline Tween 20): 20 mM of Tris-base 7.5 P^H^, 137 mM of sodium chloride, and 0.1% of Tween-20. To wash off the residual milk solution, membranes were washed three times for 10 minutes each in TBS-T prior to incubation in primary antibodies prepared in 3% of BSA diluted with TBST. Membranes were incubated in primary antibody on a rocker at 4°C overnight. Membranes were washed three times for 10 minutes each in TBST before incubating in horse radish peroxidase conjugated secondary antibodies for 1 h at room temperature. After membranes were washed three times for 10 minutes each in TBST, antibody binding detection was performed using enhanced chemiluminescence (ECL) horseradish peroxidase substrate detection kit (Millipore, Hertfordshire, UK). G: BOX Chemi-XR5 (Syngene, Cambridgeshire, UK) was used to acquire images.

### Protein synthesis (SUnSET)

Measurement of the protein synthesis was done using SUnSET (Surface sensing of translation) technique, as was previously described [44], except that puromycin was incorporated for 24 h. Briefly, fully formed myotubes were pre-treated with 1 µM of puromycin (P8833, Sigma-Aldrich) simultaneously with the indicated treatments for 24 h in a serum free media. Subsequently, protein synthesis was stimulated by 100 nM of insulin for 30 minutes in a serum-free media prior to cell lysis. The incorporation of puromycin-labelled peptides was determined by western blotting with anti-puromycin antibody (Sigma-Aldrich P8833; 1:1000).

### Glucose uptake measurements

Glucose uptake was measured using the glucose uptake ‘Promega Glo™ assay kit’. Briefly, 20,000 cells were seeded into each well of a 96-well plate. At confluent, differentiation was initiated by adding 100 µl DMEM + 2% horse serum. C2C12 WT and MuRF1 KO myotubes pre-treated with 1 µM of dexamethasone (Dex) or 0.1% ethanol (vehicle control) for 24 hours were serum starved overnight before the assay. After overnight serum starvation, wells were once washed with 100 μl DPBS and myotubes were incubated with or without 1 µM insulin under a 5% CO_2_ incubator at 37°C for 1 hour. After the incubation, myotubes were once washed with 100 μl DPBS. Next, 50 μl 1 mM 2DG in DPBS was added to each well (including myotube free wells as blank control) for 30 min at 25°C. Twenty-five μl of Stop Buffer was added with a brief shaken to lyse cells at 25°C. Next, samples were transferred to 384 well plate at 15 ul/well. Five µl of Neutralization Buffer was then applied to each well with brief shake. Finally, 20 μl of the 2DG detecting reagent was then applied to all wells with brief shake before dark-adapted for one hour at 25°C. Retained luminescence was then acquired using a BMG Labtech FLUOstar Omega microplate reader (Aylesbury, UK). Control wells (2DG: without myotubes), provided the assay background and were subtracted from all conditions.

## Data Analysis

ImageJ/Fiji (National Institutes of Health, USA) was used for protein band quantification. The proteins of interest were normalised in relation to the experimental controls.

## Statistical analysis

GraphPad Prism Software version 9 (San Diego, California, USA) was used to run all the statistical analysis. One-way analysis of variance was used for multiple group comparisons. For time course and dose response experiments, one-way analysis of variance was performed before Dunnett’s multiple comparisons post-hoc test analysis. Two-way analysis of variance of genotype (WT vs KO) and treatment (Con vs DEX) was performed before Tukey’s multiple comparisons post-hoc test analysis. All presented data are mean ± SD. Statistical significance was set at P < 0.05. The number of biological replicates and independent experiments performed is described in the figure legends.

## Data Availability

The data presented in this study are available on request from the corresponding author.

## Abbreviations

Akt: protein kinase B
ANOVA: analysis of variance
ATP: Adenosine triphosphate
BSA: bovine serum albumin
Cas9: CRISPR-associated protein
CO_2_: carbon dioxide
Con: control
CRISPR: clustered regularly interspaced short palindromic repeats
Dex: Dexamethasone
DMEM: Dulbecco’s modified eagles’ medium
DNA: deoxyribonucleic acid
ER: endoplasmic reticulum.
FoxO: forkhead box
GAPDH: glyceraldehyde 3-phosphate dehydrogenase
GFP: green fluorescent protein
gRNA: Guide RNAs
GSK-3: Glycogen synthase kinase-3
h: hours
HCl: hydrochloric acid
HECT-type: homologous to the E6-AP C-terminus
IBR: Institute of Biomedical Research
IGF-1: insulin like growth factor 1
IRS1: insulin receptor substrate 1
kDa: kilodalton
KI: knock in
KO: knock out
mg: milligram
ml: millilitre
mmol: millimolar
MPB: muscle protein breakdown
MPS: muscle protein synthesis
mRNA: messenger ribonucleic acid
mTOR: mammalian target of rapamycin
mTORC1: mammalian target of rapamycin complex 1
mTORC2: mammalian target of rapamycin complex 2
MuRF1: muscle-specific ring finger 1
MuRF2: muscle-specific ring finger 2
MuRF3: muscle-specific ring finger 3
MyBP-C: myosin-binding protein C
MyLC1: Myosin Light Chain 1
MyLC2: Myosin Light Chain 2
O_2_: oxygen
PBS: phosphate buffered saline
pH: potential of hydrogen
PI3K: phosphoinositide-3-kinase
PVDF: polyvinylidene fluoride
RhTRIM72: Human recombinant TRIM72
RING: Really Interesting New Genes
RNA: ribonucleic acid
SD: standard deviation of the mean
SDS: sodium dodecyl sulphate
SDS-PAGE: sodium dodecyl sulfate–polyacrylamide gel electrophoresis
SEM: standard error of the mean
Ser: serine
TBST: tris-buffered saline Tween-20
Thr: threonine
TRIM25: Tripartite Motif Containing 25
TRIM32: Tripartite Motif Containing 32
TRIM63: Tripartite Motif Containing 63
TRIM72: Tripartite Motif Containing 72
UPS: ubiquitin proteasome system
WT: wild type

## Acknowledgements

We would like to thank Dr Jonathan Barlow for assisting with glucose uptake data.

## Funding info

I.M. was supported by a Nigeria Tertiary Education Trust Fund Oversea Scholarship (TETF/ES/UNIV/KOGI/ASTD/2018). Y.N. was supported by the Postgraduate Research Scholarship Fund at the University of Birmingham.

## Author contributions

I.M. conceived and designed research; I.M., performed experiments; I.M., A.P.S., and Y.N., analyzed data; I.M., interpreted results of experiments; I.M. and Y.N. prepared figures; I.M. drafted manuscript; I.M., Y.N., and A.P.S. edited and revised manuscript; I.M., Y.N., and A.P.S, approved final version of manuscript.

## Conflict of interest

The authors declare that they have no conflict of interest

## Notes

### Competing Interest Statement

The authors have declared no competing interest.

